# Conserved inter-domain interactions drive trans-Golgi network localisation and trafficking of homologous copper-ATPases

**DOI:** 10.1101/2025.05.01.651363

**Authors:** Sharon Mary Jose, Saptarshi Maji, Mrittika Paul, Aayatti Mallick Gupta, Arnab Gupta

## Abstract

Polytopic transmembrane Copper-ATPases are central regulators of the essential, redox-active micronutrient copper in all organisms. The vertebrate Cu-ATPase homologs ATP7A and ATP7B facilitate excess copper efflux, in addition to metalating cuproproteins. ATP7A and ATP7B exhibit copper-responsive basolateral and apical trafficking polarity respectively, in polarized epithelia. To elucidate the role of conserved functional domains in copper-responsive trafficking and copper transport, we substituted the copper-binding amino-terminal, Nucleotide-Binding (NBD) and C-terminal domain of ATP7B with that of ATP7A either singly or in combinations. The six chimeric Cu-ATPases generated, although functionally active, exhibited altered trafficking phenotypes. Notably, the N-terminal-substituted ATP7B exhibited constitutive trafficking to the basolateral membrane; however, the N-terminal and NBD double-substituted chimera gained steady-state TGN localization, indicating a putative interaction between the two domains critical for its TGN-localization. We analyzed orthologous Cu-ATPase domain sequences from diverse organisms and found that the N-terminal domain and NBD showed similar evolutionary relationships, unlike that of C-terminal domain, suggesting their co-evolution. This study for the first time correlates evolutionary stringency imparted onto Cu-ATPase domains with their significance in proper trafficking regulation.

## Introduction

Copper is an essential micronutrient (Mustafa & Alsharif, 2018; Olivares & Uauy, 1996). However, copper levels in all cells need to be tightly regulated, given the high redox potential of this metal. Although copper is involved in crucial physiological processes like cellular respiration (Wu et al., 2024), iron metabolism (Ruiz et al., 2021), neuroendocrine function (Eipper et al., 1983), host defense against microbial pathogens (Fu et al., 2014; Paul et al., 2022,2024), free copper ions can participate in Fenton-like reactions to generate potentially toxic reactive oxygen species (Zischka and Lichtmannegger, 2019) as well as displace other metal ions from metalloenzymes (Gaetke & Chow, 2003; Macomber & Imlay, 2009). Copper-transporting P-type ATPases (Cu-ATPases), found in all life forms are central regulators of cellular copper (La Fontaine & Mercer, 2007; Smith et al., 2014). Cu-ATPases evolved from being static cell membrane transporters in prokaryotes to dynamic trafficking entities in multicellular eukaryotes (Gupta and Lutsenko; 2012). The intracellular location of the Cu-ATPases in multicellular eukaryotic cells is dictated primarily by the intracellular copper concentration (La Fontaine & Mercer, 2007; Lutsenko et al., 2007; Voskoboinik & Camakaris, 2002). Under high cellular copper conditions, they traffic out of trans-Golgi network (TGN) to lysosomes or cell membrane to sequester and eliminate excess copper. When copper levels subsequently recede, these Cu-ATPases recycle to the TGN (Petris & Mercer, 1999; Cater et al., 2006).

Vertebrate organisms possess two homologous Cu-ATPases, i.e., Cu-ATPase-1 or ATP7A and Cu-ATPase-2 or ATP7B; interestingly, the two Cu-ATPases show distinct trafficking polarity in response to high copper in epithelial cells (Greenough et al., 2004; Nyasae et al., 2014). ATP7A, which is expressed in all cell types except hepatocytes, traffics to the basolateral membrane in polarized epithelial cells. In enterocytes, ATP7A facilitates the absorption of dietary copper by transporting it across the basolateral membrane, which interfaces with the bloodstream, thereby enabling systemic copper distribution (Nyasae et al., 2007). Mutations in human *ATP7A* result in Menkes disease, a severe systemic copper deficiency disorder (Tümer & Møller, 2010). On the other hand, ATP7B is expressed abundantly in hepatocytes, and co-expressed with ATP7A in kidney and brain (Fanni et al., 2009; Linz et al., 2008; Telianidis et al., 2013). In adult hepatocytes, where it is the sole Cu-ATPase, ATP7B traffics to the apical membrane to pump out the excess copper into the bile canaliculus, from where copper is eventually excreted out of the body via feces (Roelofsen et al., 2000). Mutations in human *ATP7B* lead to Wilson disease, where excess copper accumulates and damages the liver, kidney, and brain (Bull et al., 1993).

All Cu-ATPases possess multiple cytosolic and transmembrane (TM) domains with signature P-type ATPase motifs that enable copper transport across membranes. Besides eight TM domains, vertebrate Cu-ATPases possess the following cytosolic domains– (1) N-terminal domain harboring six copper-binding sites, each site containing an MXCXXC motif. ATP7B possesses an extra 63 aa region which has signals for its TGN-localization and apical targeting in polarized hepatocytes (Braiterman et al., 2008) (2) Actuator domain responsible for dephosphorylation, present between TM4 and TM5, (3) nucleotide-binding (N) and phosphorylation (P) domains, responsible for ATP-binding via SEHPL motif and hydrolysis via DKTG motif, respectively, present in the loop between TM6 and TM7 and (4) C-terminal domain which harbours conserved dileucine motif, required for copper-dependent trafficking, TGN localization and retrograde trafficking (Petris & Mercer, 1999; Zhu,S et al., 2016; Greenough et al., 2004; Ruturaj et al., 2024).

Trafficking regulators like Adaptor protein complexes like AP1 (Cancino et al., 2007; H Fölsch et al., 1999; Gan et al., 2002; Gonzalez & Rodriguez-Boulan, 2009; Gravotta et al., 2007, 2012; Ruturaj et al., 2024; Jain et al., 2015), Clathrin (Deborde et al., 2008; Holloway et al., 2013), AIPP1 (Stephenson et al., 2005), microtubule (Jaulin et al., 2007; Perez Bay et al., 2013; Weisz & Rodriguez-Boulan, 2009) and actin regulatory proteins (Braiterman et al., 2009; Gupta et al., 2016; Salvarezza et al., 2009) have been reported to play roles in the trafficking of ATP7A and ATP7B. Mutational studies have also deciphered the roles of a few motifs in the N-terminal and C-terminal domains of ATP7A (Zhu, S. et al., 2016; Veldhuis, N. A. et al., 2009; Francis M. et al., 1999; Goodyer I. et al., 1999) and ATP7B (Braiterman et al., 2015; Cater et al., 2004; Hasan & Gupta, 2012; Das S. et al., 2022; Ruturaj et al., 2024) in mediating steady-state TGN localization and copper-responsive trafficking. Interestingly, a few *in silico* (Oradd F. et al., 2022) and *in vitro* structural studies (Yang et al., 2023; Bitter R. et al., 2022; Hasan & Gupta, 2012; LeShan E. et al., 2010; Braiterman et al., 2015) have postulated the significance of inter-domain interactions, involving the cytosolic ATP7B domains in copper transport and trafficking. Similar studies in ATP7A are limited though (Banci et al., 2009). Since ATP7A and ATP7B branched out from a single Cu-ATPase during the evolution of vertebrates from lower chordates, we asked whether the cytosolic domains between these transporters could be exchanged without disrupting the trafficking and transport functions of the proteins. We aimed to understand the inter-domain interactions that play roles in their copper-transport activity and copper-responsive localization in epithelial cells. To this end, we considered three cytosolic domains of ATP7B that have been reported to be important in trafficking – N-terminal domain, NBD (consisting of the N- and P-domains), and C-terminal domain, and replaced these domains with those of its homolog ATP7A, singly and in pairs, and tested the copper-transport activity and trafficking ability of the resulting chimeric Cu-ATPases. To better interpret the observations, we dissected the evolutionary patterns of these key cytosolic domains across diverse life forms. Our results provide insights into the extent of divergence of the key Cu-ATPase domains and the significance of their interdomain interactions in regulating the intracellular localization and activity of the Cu-ATPases.

## Results

### ATP7A and ATP7B show distinctive copper-dependent localization in epithelial cells

Previous studies from our group have shown that at basal copper levels, ATP7A and ATP7B reside at the TGN in different sub-domains. At high copper conditions, ATP7A traffics to basolateral membrane directly, and ATP7B traverses distinct endosomal compartments *en-route* to the apical membrane, in polarized epithelial cell model MDCK cells (Ruturaj et al., 2024). These cells of canine origin endogenously express ATP7A and ATP7B, but since antibodies against dog ATP7A and ATP7B are commercially unavailable, we transiently transfected MDCK cells with fluorescently tagged ATP7A (mKO2-HA-ATP7A) or ATP7B (EGFP-ATP7B) and exposed the cells to three conditions – basal, high copper (one-hour 75µM CuCl_2_), or copper-depletion post-copper treated - to activate recycling back to TGN (one-hour 75µM CuCl_2_ followed by two hours’ treatment with 100µM Bathocuproinedisulfonic acid (BCS), a cell-impermeable Cu-chelator). We verified the localization of these wild type Cu-ATPases during unpolarized and polarized MDCK cell stages, at the three different copper conditions, to detect differences in trafficking regulation that occur during epithelial cell polarization.

Under basal copper condition, mKO2-HA-ATP7A was localized in the trans-Golgi Network (TGN), marked by Golgin-97, in both unpolarized **(Fig. 1A)** and polarized **(Fig. 1B)** MDCK cells. Upon elevation in intracellular copper levels (+Cu), in unpolarized cells, ATP7A traffics out of TGN and localized to the cell membrane (marked by cortical actin network), whereas, in polarized MDCK cells, ATP7A trafficked to the basolateral membrane. Following copper chelation (+Cu to -Cu), ATP7A was seen to recycle back to the TGN. Fraction of mKO2-HA-ATP7A in TGN and in cell membrane/basolateral membrane was quantified using Manders Overlap Coefficient formula and is depicted in **Fig. 1C** (unpolarized MDCK) and **Fig. 1D** (polarized MDCK).

**Fig. 1.**
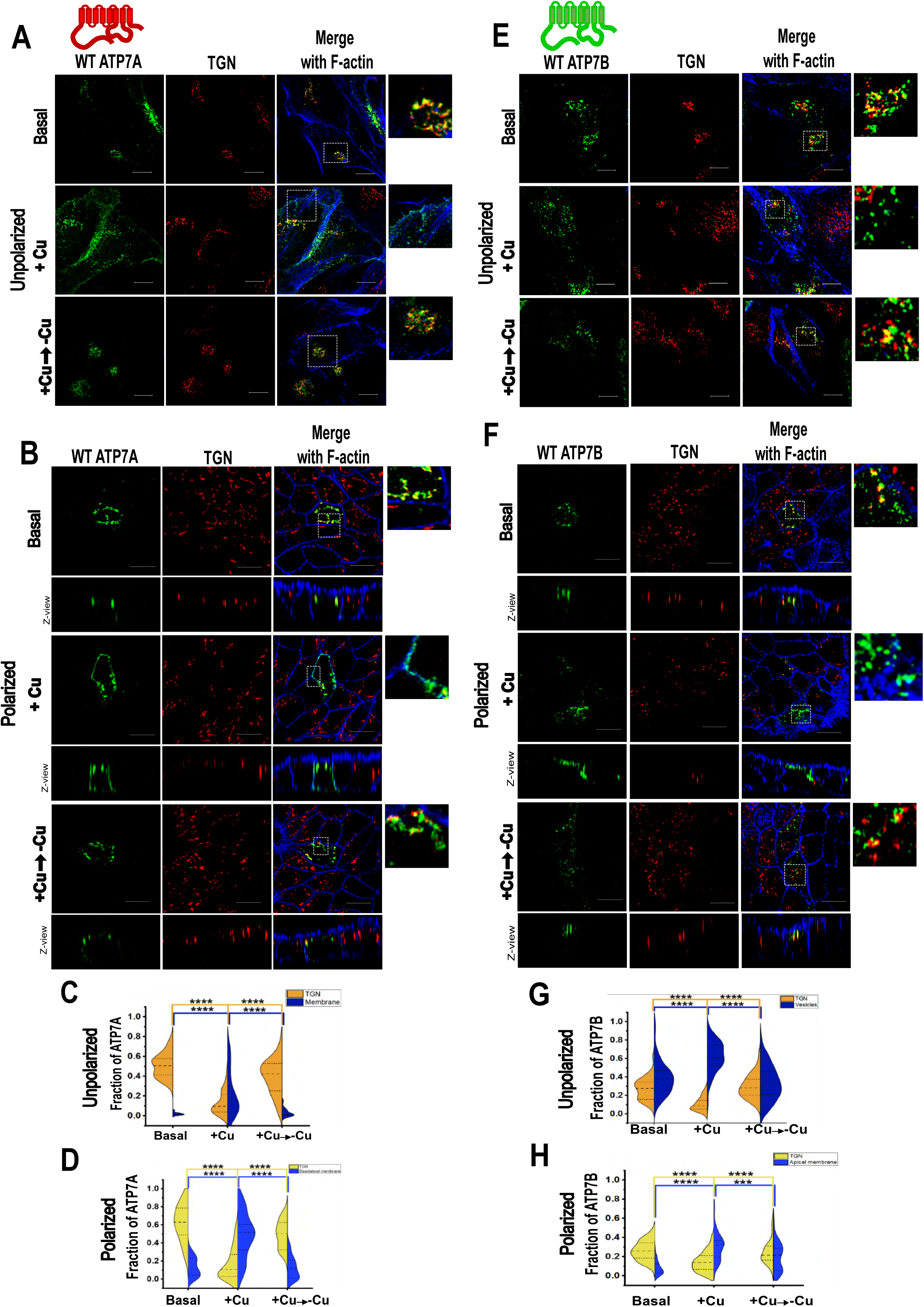
Localization of homologous copper ATPases at different copper concentrations in unpolarized and polarized MDCK cells. (A) Unpolarized and (B) polarized MDCK cells transfected with mKO2-HA-ATP7A under basal, copper-treated and copper depletion post-copper treated conditions. TGN is marked by Golgin-97. ATP7A, in high copper conditions, traffics from the TGN to cell membrane/basolateral membrane in polarized cells and returns to the TGN upon chelation of excess copper. (C) and (D) depicts graphically the fraction of the protein (Manders Overlap Coefficient) present in TGN and cell membrane/basolateral membrane (marked by ATP1A) under the three copper conditions tested for unpolarized and polarized cells respectively. Sample size N=52, 41 and 55 (unpolarized) and N=37, 52 and 30 (polarized) for basal, Cu-treated and +Cu to -Cu conditions, respectively. (E) Unpolarized and (F) polarized MDCK cells transfected with EGFP-ATP7B under the same three copper conditions are shown. ATP7B traffics from the TGN under high copper conditions, to post-TGN compartments (not marked here) in unpolarized MDCK cells, or to the apical membrane in polarized cells, and recycles back to TGN when copper levels decrease. (G) and (H) show plots quantifying the fraction (Manders Overlap Coefficient) of EGFP-ATP7B residing in TGN and post-TGN compartments (unpolarized cells) or the apical membrane (marked by gp135) in polarized cells, at the three copper levels tested. Sample size N =48, 42 and 52 cells (unpolarized cells) and N = 61, 75 and 43 (polarized cells) for basal, Cu-treated and +Cu to - Cu conditions, respectively. Scale bar: 10µm. (* copper treatment for all experiments: 75µM CuCl_2_ for one hour and +Cu to -Cu: 75µM CuCl_2_ for one hour, followed by copper chelation by 100 µM BCS for two hours).

Similarly, EGFP-ATP7B, at basal conditions, was found to be primarily localized in the TGN and a small fraction in post-TGN vesicles in unpolarized MDCK cells **(Fig. 1E)**. ATP7B is known to constitutively recycle between TGN and recycling endosomes (Maji et al., 2023; Ruturaj et al., 2024). Upon copper treatment (+Cu), ATP7B was seen to be present in post-TGN compartments, and upon further copper depletion, majority of the ATP7B pool recycled back to the TGN. In polarized cells **(Fig. 1F)**, at basal condition, ATP7B was present at the TGN and post-TGN compartments and upon copper treatment, majority of the protein trafficked to the apical membrane. When the excess copper was chelated out with copper-chelator, ATP7B recycled back to the TGN. Fraction of ATP7B in TGN and in post-TGN compartment/apical membrane was quantified using Manders Overlap Coefficient formula and is depicted in **Fig. 1G** (unpolarized MDCK) and **Fig. 1H** (polarized MDCK).

### Orthologous vertebrate Cu-ATPase domains show comparable sequence similarities

Both Cu-ATPase homologs display differential apico-basolateral trafficking in polarized cells despite sharing about 55% sequence identity. To gain insights into the domain-specific differences behind these phenotypes, we analysed three cytosolic domains constituting these Cu-ATPases ̶ the N-terminal domain, NBD loop (comprising of the N- and P-domains) and the C-terminal domain ̶ that have been previously implicated in trafficking, as mentioned earlier. Since vertebrates were the first animals to harbour both ATP7A and ATP7B, we retrieved protein sequences of these Cu-ATPase domains from eight different vertebrate species including humans, viz. sea lamprey, frog, green lizard, shark, zebrafish, chicken, and rat, from NCBI database and aligned them using Clustal Omega software. **Fig. 2A** depicts a segmented bar graph showing the sequence identities of the homologous Cu-ATPase domains in eight different vertebrate species. The C-terminal domain showed the least homology (average ∼37%), followed by the N-terminal domain (average ∼44%), whereas the NBD of both Cu-ATPases shared the highest degree of homology (average ∼67%). Interestingly, the homologous domains were of comparable similarities across the different vertebrate taxa. Key differences in the N-terminal, N-domain and C-terminal domains have been highlighted in **Fig. 2B** To elucidate how these domain(s) influence trafficking and copper transport as well as to understand their functional divergence, we exchanged the three cytosolic domains of ATP7B one by one, with those from its homologue ATP7A.

**Fig. 2.**
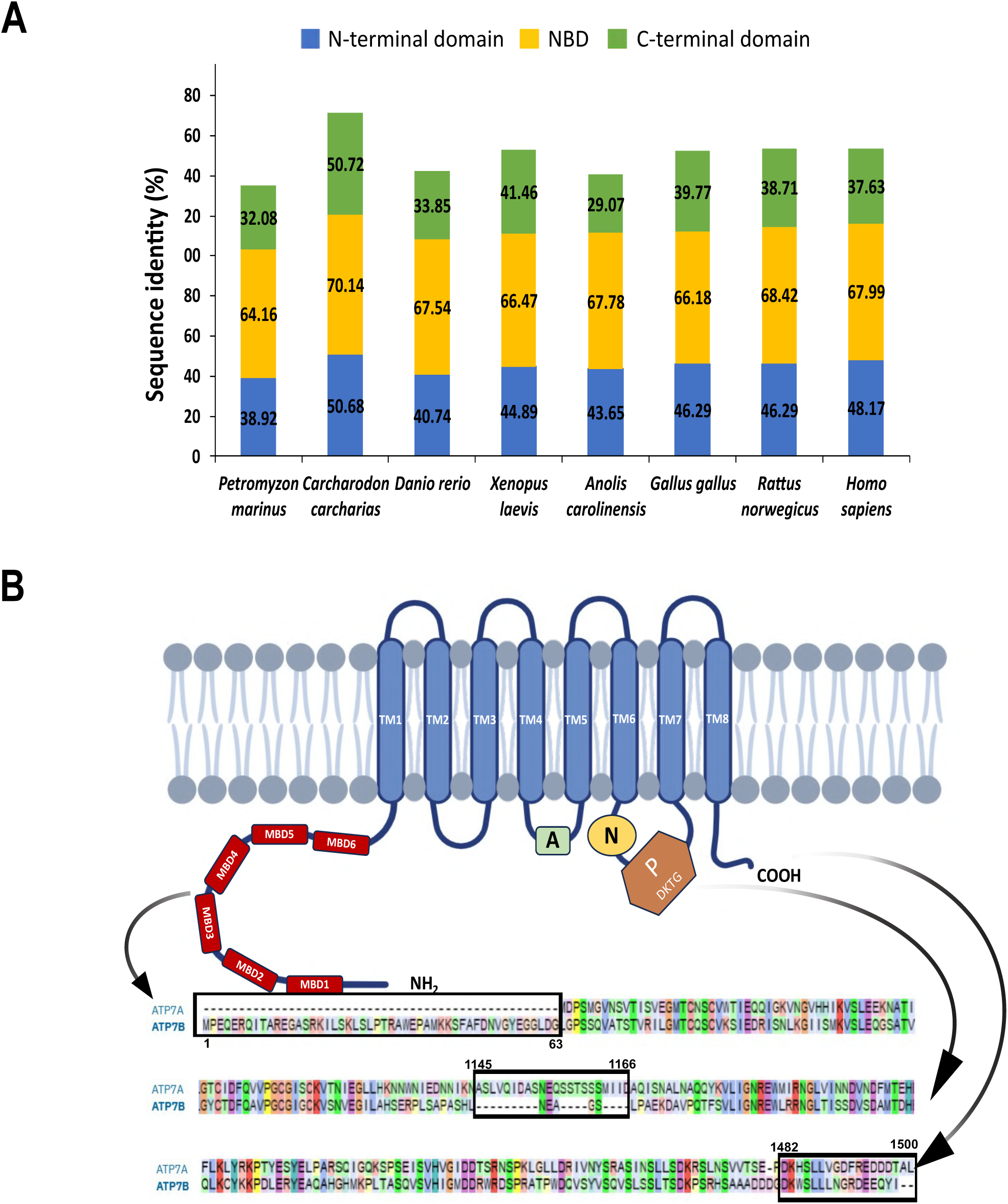
Graph showing sequence identity between Cu-ATPase-1 and CuATPase-2 domains of different vertebrates and key domain differences in human Cu-ATPases. (A) Amino acid sequences of N-terminal, Nucleotide binding and C-terminal domains of homologous copper ATPases of eight vertebrates were aligned and percentage identities were plotted as segmented bar graphs. All eight vertebrate Cu-ATPases show comparable homology between the three domains. (B) Cartoon of a Cu-ATPase highlighting the key sequence differences between the N-terminal, NBD and C-terminal domains of human Cu-ATPases ATP7A and ATP7B.

### Substitution of ATP7B N-terminal domain with that of ATP7A results in basolateral targeting of the chimera

It has been previously reported that the distal end of N-terminus of ATP7B has extra 63 residues, that is absent in ATP7A. A nine residue stretch F^37^AFDNVGYE^45^ within these 63 amino acids was implicated in proper TGN-localization and apical targeting in polarized hepatocytes. Deleting this region leads to its mistargeting to the basolateral membrane (Braiterman et al., 2008). No such comparable motif has been detected in the N-terminal domain of ATP7A though.

At the outset, we replaced the N-terminal domain of ATP7B with that of ATP7A – the resultant chimeric construct obtained by Hi-Fi cloning ̶ EGFP-(7A-NT)-ATP7B - was sequenced and after confirmation of sequence, MDCK cells were transfected with the construct and exposed to identical copper conditions that were used for wild type Cu-ATPases in results section 1. In unpolarized MDCK cells (**Fig. 3A**), at basal conditions, this chimera was found to be present in the TGN, vesicularized compartments, as well as on the cell membrane. In polarized cells too, the chimera exhibited similar phenotype (**Fig. 3B**). When these cells were exposed to high copper conditions, TGN-localized and vesicularized (7A-NT)-ATP7B trafficked to the cell membrane in unpolarized and basolateral membrane in polarized cells. Interestingly, when this excess copper was chelated out with BCS, the chimera was retained at the basolateral membrane and failed to recycle to the TGN. Remarkably, vesicularized (7A-NT)-ATP7B population recycled back to the TGN as seen by a significant increase in the TGN localization of these chimera in the +Cu to -Cu condition. Fraction of EGFP-(7A-NT)-ATP7B in TGN and cell membrane/basolateral membrane, quantified using Manders Overlap Coefficient formula is shown in **Fig. 3C** (unpolarized MDCK) and **Fig. 3D** (polarized MDCK). (7A-NT)-ATP7B showed constitutive trafficking to the cell membrane. This finding highlights the importance of N-terminal domain of ATP7B in copper-dependent localization of the protein at the TGN and correct trafficking to the apical membrane in polarized cells.

**Fig. 3.**
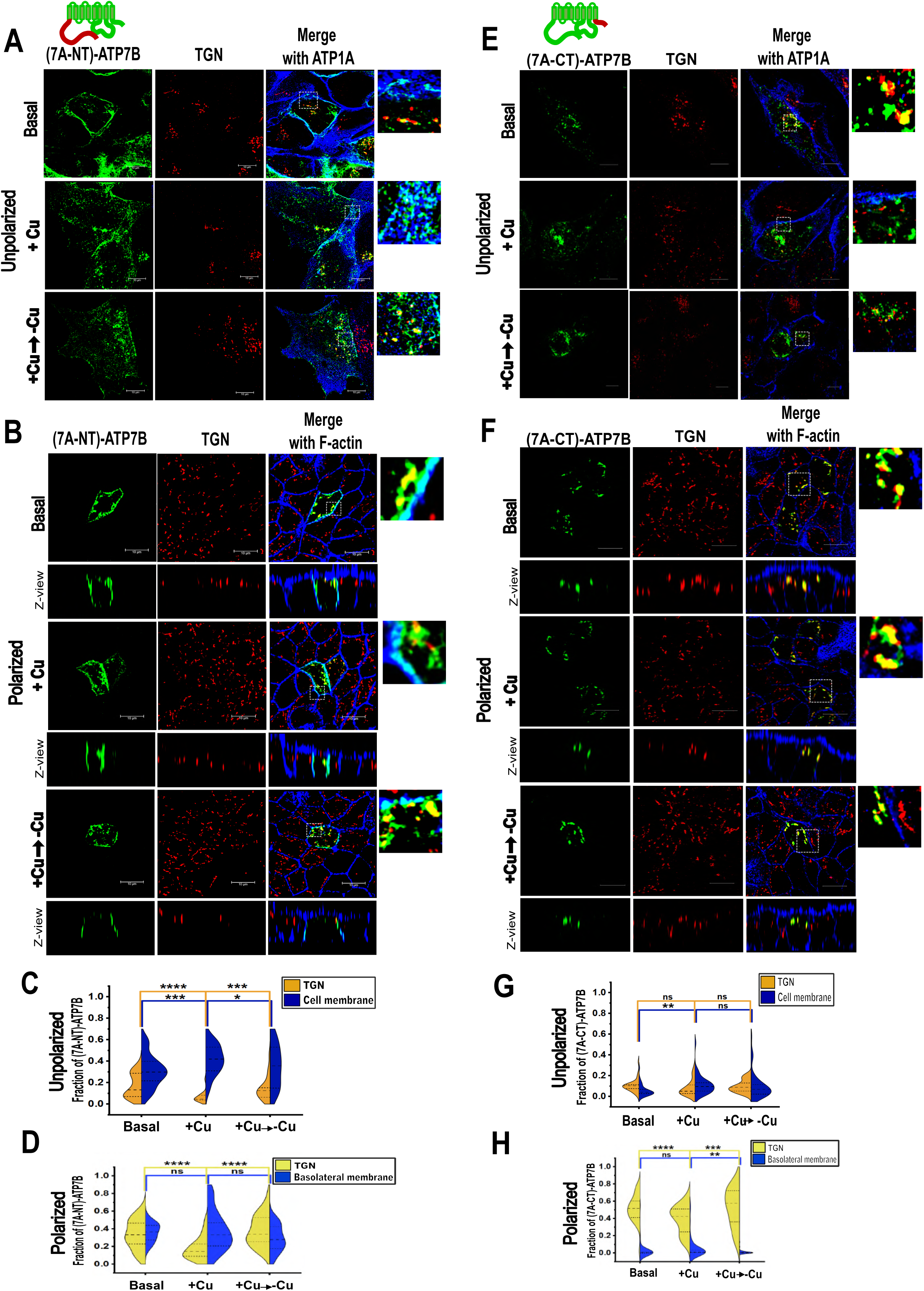
Amino-terminal, but not carboxy-terminal domain substitution of ATP7B, affects its steady-state localization at TGN. TGN is marked by Golgin-97. (A) and (B) shows the localization of EGFP-(7A-NT)-ATP7B in unpolarized and polarized MDCK cells respectively under basal, copper-treated and copper depletion post-copper treated conditions. These chimeric Cu-ATPase molecules were present in TGN, post-TGN compartments and cell membrane/basolateral membrane in all three copper conditions tested. Quantification of the fraction (Manders Overlap Coefficient) of these proteins in the TGN/membrane is shown in (C) for unpolarized MDCK and (D) for polarized MDCK. Sample size: N=31, 27, 29 for unpolarized cells and N=30, 36, 34 for polarized cells at basal, copper-treated and +Cu to -Cu conditions, respectively. Localization of the second chimera, EGFP-(7A-CT)-ATP7B in unpolarized (E) and polarized (F) MDCK cells, under the same three conditions is shown. In unpolarized stage, this chimera was seen to be present in post-TGN compartments and TGN at basal conditions and it trafficked to the cell membrane to a low extent in response to high copper, but it did not traffic back to the TGN even upon subsequent copper chelation. In polarized stage, the chimera showed predominant TGN localization at basal conditions, and trafficked out to post-TGN compartments in response to copper and recycled to TGN upon subsequent copper chelation. (G) and (H) shows the quantification (Manders Overlap Coefficient) of the colocalization of this chimera with TGN and cell membrane/basolateral membrane (ATP1A) in unpolarized (N=32, 35 and 47) and polarized cells (N= 38, 45 and 49 cells) at basal, copper-treated and +Cu to -Cu conditions, respectively. Scale bar: 10µm

For ATP7A, the dileucine motif (L^1487^-L^1488^) present in the carboxy terminal has been reported to be important for both TGN localization and basolateral trafficking in polarized MDCK cells, besides playing a role in its endocytic retrieval. Deleting this motif led to the apical mistargeting of the mutant ATP7A, regardless of copper levels (Greenough et al., 2004). It has also been showed that deleting a PDZ target ‘DTAL’ motif from the C-terminus led to the apical targeting of the mutant ATP7A under elevated copper. On the other hand, for ATP7B, the tri-leucine motif mutation leads to its constitutive cell membrane localization (Braiterman et al., 2011). Based on these findings, we constructed a second chimeric ATP7B with its C-terminal domain replaced by that of ATP7A. In unpolarized cells **Fig. 3E**, EGFP-(7A-CT)-ATP7B was found to be majorly localized in vesicular compartments, and to a lower extent at the TGN, at all three tested copper conditions. However, when cells were polarized **Fig. 3F**, (7A-CT)-ATP7B was seen to be largely present in the TGN at basal conditions, and in response to high copper, the chimera trafficked out of the TGN in vesicular compartments, but did not reach the cell membrane. Further, upon treatment with copper chelator, the chimera recycled back to the TGN. **Fig. 3G** shows the fraction of this chimera in TGN/cell membrane in unpolarized MDCK cells, while **Fig. 3H** shows the fraction of this chimera in TGN/basolateral membrane in polarized cells. These results indicated that substituting the C-terminal domain of ATP7B with that of ATP7A, affected its steady-state TGN-localization in unpolarized cells. In polarized cells, not only was this phenomenon largely rescued, the chimera could also traffic out and into the TGN in response to copper. It is possible that the presence of the conserved dileucine motif in the two homologous domains caused the chimera to show normal Cu-responsive trafficking phenotype, at least in polarized MDCK.

We then generated a double-substituted chimeric ATP7B by replacing both its N- and C-termini with those of ATP7A. The resultant chimeric Cu-ATPase ̶ EGFP-(7A-NT, CT)-ATP7B– in unpolarized cells **(Fig. 4A)** was found to be present at post-TGN compartments majorly and to a lower extent at the TGN at basal conditions. When subjected to high copper levels, a pool of this chimera could traffic to the cell membrane, but even upon subsequent copper-chelation, there was no change in the chimera pool at the TGN or the cell membrane. In polarized cells **(Fig. 4B),** at basal conditions, the chimera showed a constitutive targeting to the basolateral membrane, besides being at the TGN and post-TGN compartments. There was no change in its localization even upon copper addition. Remarkably, upon subsequent copper chelation, there was endocytosis of the membrane pool of the chimeric protein, even though they failed to return to the TGN. The same has been quantified in **Fig. 4C** and **4D**. This indicated that this doubly-substituted chimera did not traffic constitutively to the cell membrane in unpolarized cells, as opposed to (7A-NT)-ATP7B. But in polarized cells, the chimera showed the same phenotype as (7A-NT)-ATP7B, except that this second chimera could be endocytosed from the basolateral membrane upon copper chelation, although it could not recycle back to the TGN.

**Fig. 4.**
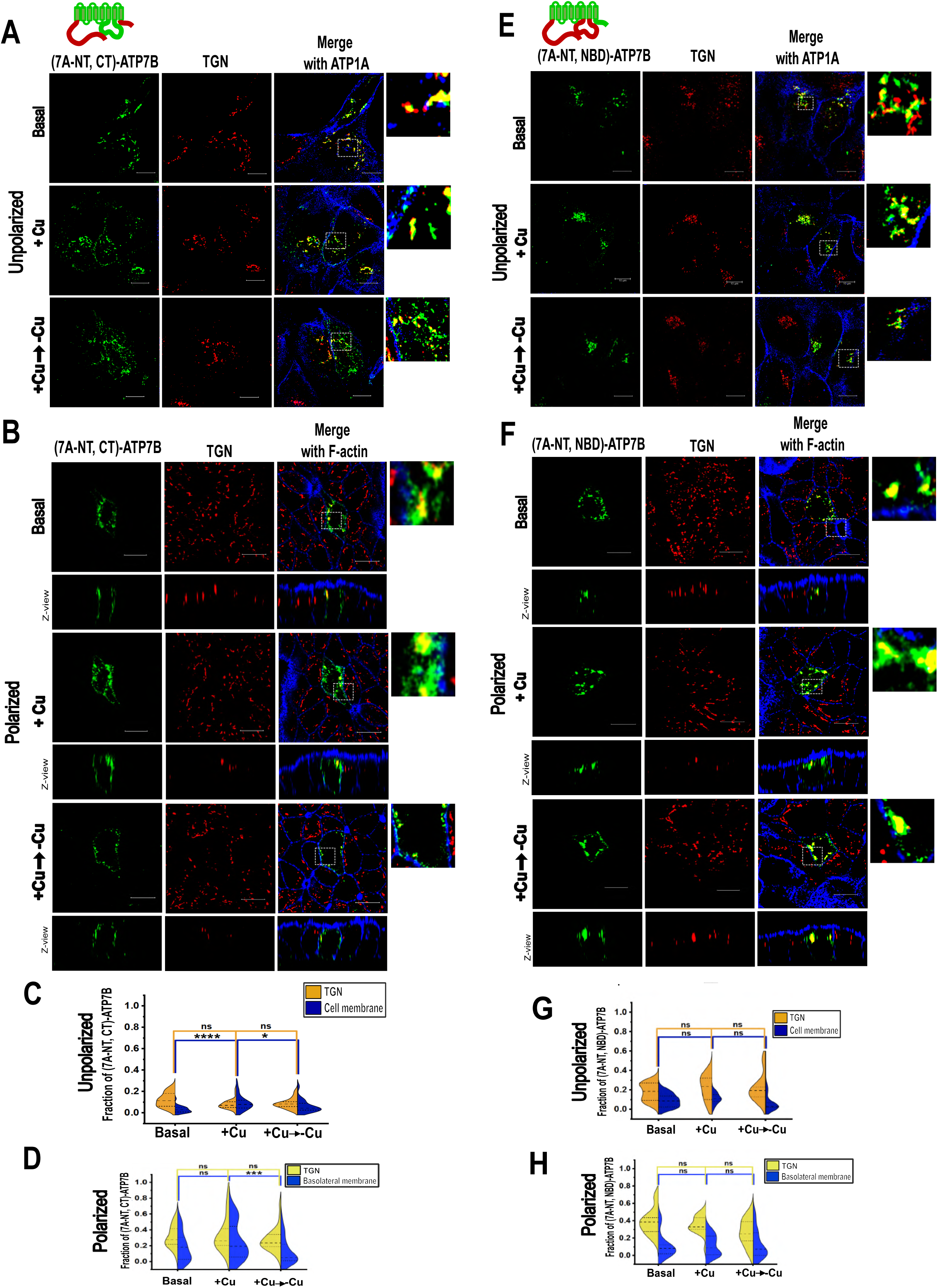
Substitution of NBD, but not C-terminal domain, along with N-terminal domain of ATP7A in ATP7B, stabilized TGN-retention in polarized MDCK. TGN is marked by Golgin-97. (A) Unpolarized and (B) polarized MDCK cells transfected with EGFP-(7A-NT, CT)-ATP7B at basal, copper-treated and copper depletion post-copper treated conditions are shown. This chimera trafficked to the cell membrane to an extent in response to copper in unpolarized cells, but in polarized cells, it showed constitutive trafficking to the basolateral membrane, even at basal conditions. Recycling from the membrane (but not up to the TGN) upon subsequent copper chelation was observed in polarized cells, but not in unpolarized cells. Quantification of the fraction (Manders Overlap Coefficient) of these proteins in the TGN/membrane is shown in (C) for unpolarized cells (N=31, 29 and 40 cells) and in (D) for polarized cells (N=24, 29 and 28 cells) for basal, copper-treated and +Cu to -Cu conditions. The localization of the fourth chimera, EGFP-tagged (7A-NT, NBD)-ATP7B is showed in Fig. 4E (unpolarized MDCK) and Fig. 4F (polarized MDCK) at the same three copper conditions. This chimera was found to be localized at the TGN mostly, in all copper conditions tested, at both stages of MDCK cell polarization. (G) and (H) shows the quantification of the colocalization of this chimera with TGN and cell membrane (ATP1A) in unpolarized (N=33, 32 and 36) and polarized cells (N= 41, 30 and 31 cells) at basal, copper-treated and +Cu to -Cu conditions, respectively. Scale bar: 10µm

### Complex interdomain interactions involving Nucleotide-binding domain are crucial for copper-responsive TGN localization and exit

ATP-binding loops of the two human Cu-ATPases have been shown to possess comparable affinities to adenosine nucleotides AMP, ADP and ATP (Morgan et al., 2004). But, the N-domains of the homologs possess variable unfolded loops of different lengths (H^1115^-D^1138^ in ATP7B, H^1131^-A^1175^ in ATP7A) that have been hypothesized to be important for trafficking regulation. In ATP7B, it has been hypothesized that an interaction between the N-domain and the copper-binding amino-terminal domain may be essential for TGN localization and copper-responsive trafficking (Tsivkovskii et al., 2001; Hasan and Gupta, 2012). Since, (7A-NT)-ATP7B was trafficked to the cell membrane constitutively, we wondered about the implication of additionally substituting the NBD of ATP7A in this chimera. We therefore constructed EGFP-(7A-NT-NBD)-ATP7B, and transfected MDCK cells and studied the localization phenotypes. Surprisingly, in unpolarized cells at all three copper conditions, this chimera was primarily found to be localized at TGN and post-TGN compartments, with minimal localization at the cell membrane, (**Fig. 4E**). In polarized cells (**Fig. 4F**), the chimera was largely localized at the TGN at all the three copper conditions tested. We predict that an inter-domain interaction between the N-terminal domain and the NBD of ATP7A may have caused this chimeric protein to be retained at the TGN/post TGN vesicles, unlike the (7A-NT)-ATP7B that constitutively trafficked to the cell membrane. However, this chimera could not traffic out of the TGN/post-TGN compartments in response to copper, possibly due to the existence of further complex intramolecular interactions. The same has been quantified in **Fig. 4G** (unpolarized cells) and **4H** (polarized cells).

To extend our understanding of the role of the NBD in trafficking of the copper-ATPases, we generated two other NBD-substituted ATP7B chimeras ̶ the double substituted EGFP-(7A-NBD, CT)-ATP7B and singly substituted EGFP-(7A-NBD)-ATP7B and transfected MDCK cells with these constructs. **Fig. 5A** shows the unpolarized MDCK cells transfected with a chimeric ATP7B with both NBD and C-terminal replaced by those of ATP7A. This chimeric protein, (7A-NBD, CT)-ATP7B was seen to be localized primarily at post-TGN compartments, and at the TGN to a lower extent, at basal conditions. In polarized cells (**Fig. 5B**), the chimera was seen to be present in the TGN largely and it did not traffic in response to copper. **Fig. 5C** and **D** shows the quantification of the same in unpolarized and polarized cells respectively.

**Fig. 5.**
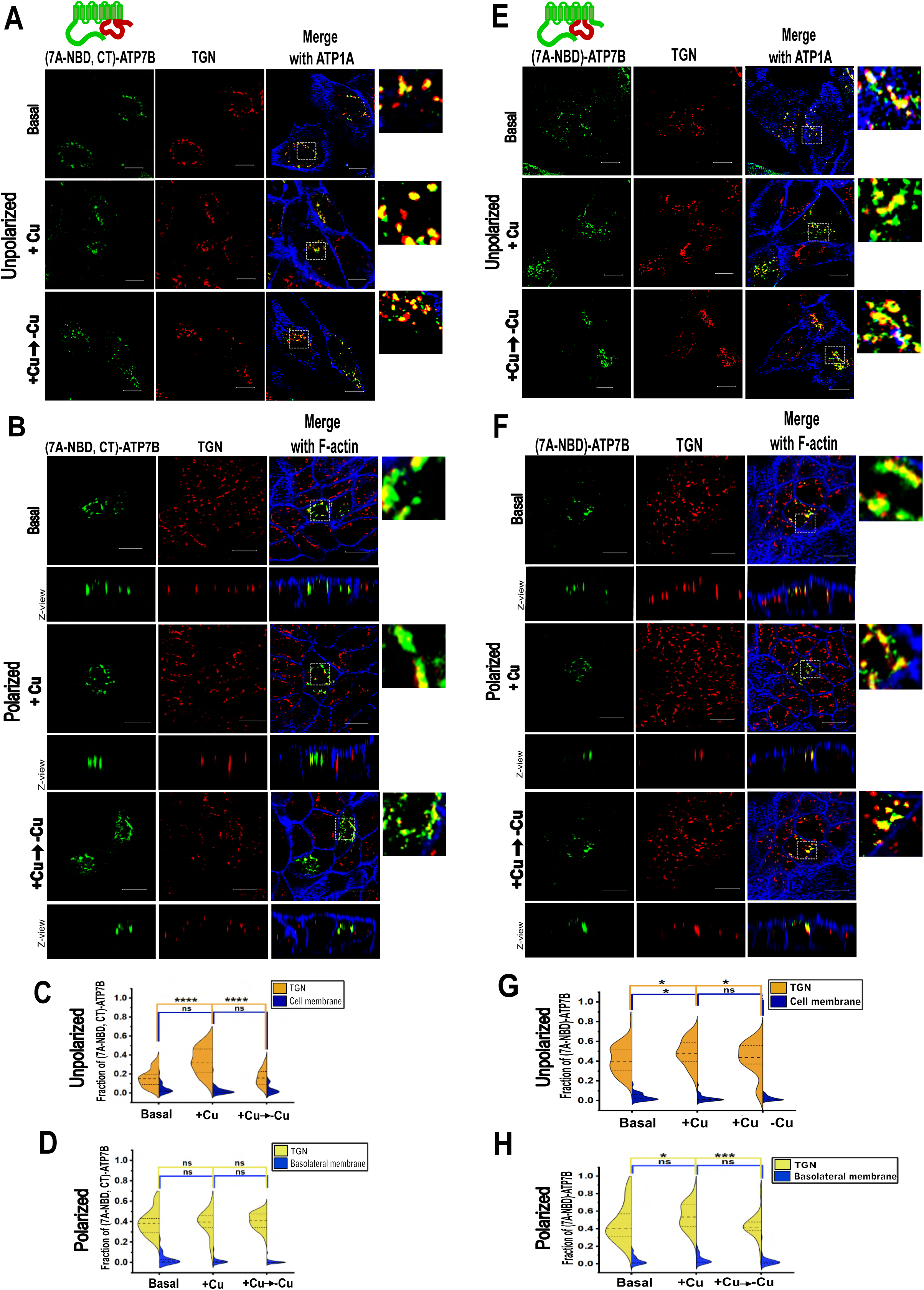
ATP7B NBD-substituted chimeras show TGN-localization but fail to show Cu-responsive TGN-exit. TGN is marked by Golgin-97. (A) and (B) shows the localization of EGFP-(7A-NBD, CT)-ATP7B in unpolarized and polarized MDCK cells, respectively, under basal, copper-treated and copper depletion post-copper treated conditions. This chimera was found to be present at post-TGN compartments and TGN in unpolarized cells, although it showed increased localization at the TGN at high copper. In polarized cells, this chimera was localized predominantly at the TGN. (C) and (D) shows the quantification of the colocalization of this chimera with Golgin-97 and cell membrane (ATP1A) in unpolarized (N=37, 35 and 25 cells) and polarized cells (N= 29, 28 and 32 cells) at basal, copper-treated and +Cu to -Cu conditions, respectively. The localization of the sixth chimera, EGFP-tagged (7A-NBD)-ATP7B is showed in Fig. 5E (unpolarized MDCK) and Fig. 5F (polarized MDCK) at the same three copper conditions. This chimera was seen to be localized at the TGN predominantly irrespective of copper levels in both stages of MDCK cell polarization. (G) shows the colocalization quantification of this chimera with Golgin-97 and membrane (ATP1A) in unpolarized (N=31, 27 and 29) and in polarized cells (N=30, 36 and 34), for basal, copper-treated and +Cu to -Cu conditions, respectively. Scale bar: 10µm

A similar phenotype was exhibited by EGFP-(7A-NBD)-ATP7B in transfected MDCK cells. The chimeric proteins were found to be present mostly at the TGN at all three copper conditions tested in both unpolarized **(Fig. 5E)** and polarized cells **(Fig. 5F). (****Fig. 5G** and **H**) shows the quantification of the same in unpolarized and polarized cells respectively. Interestingly, all the constructs with NBD of ATP7A in ATP7B failed to exhibit Cu-dependent TGN-exit, a phenotype that overrides other localization or trafficking phenotypes. This points towards the existence of complex conformational changes involving NBD and key inter-domain interactions that must be involved in facilitating trafficking of the Cu-ATPase.

### Chimeric Cu-ATPases exhibit normal copper-transport activity

**Fig. 6A** shows schematically all the chimeric constructs generated. Even though all the six chimeric constructs generated were found at the TGN, albeit to different extents, we tested their stability in comparison to the wild type ATP7A and ATP7B. We transfected HEK293T cells with plasmids encoding the chimeras, and post-24 hours expression, immunoblotting with anti-GFP was performed. Though their levels of expression slightly varied, all of them were seen to be stably synthesized in cells **(Fig. 6B).**

**Fig. 6.**
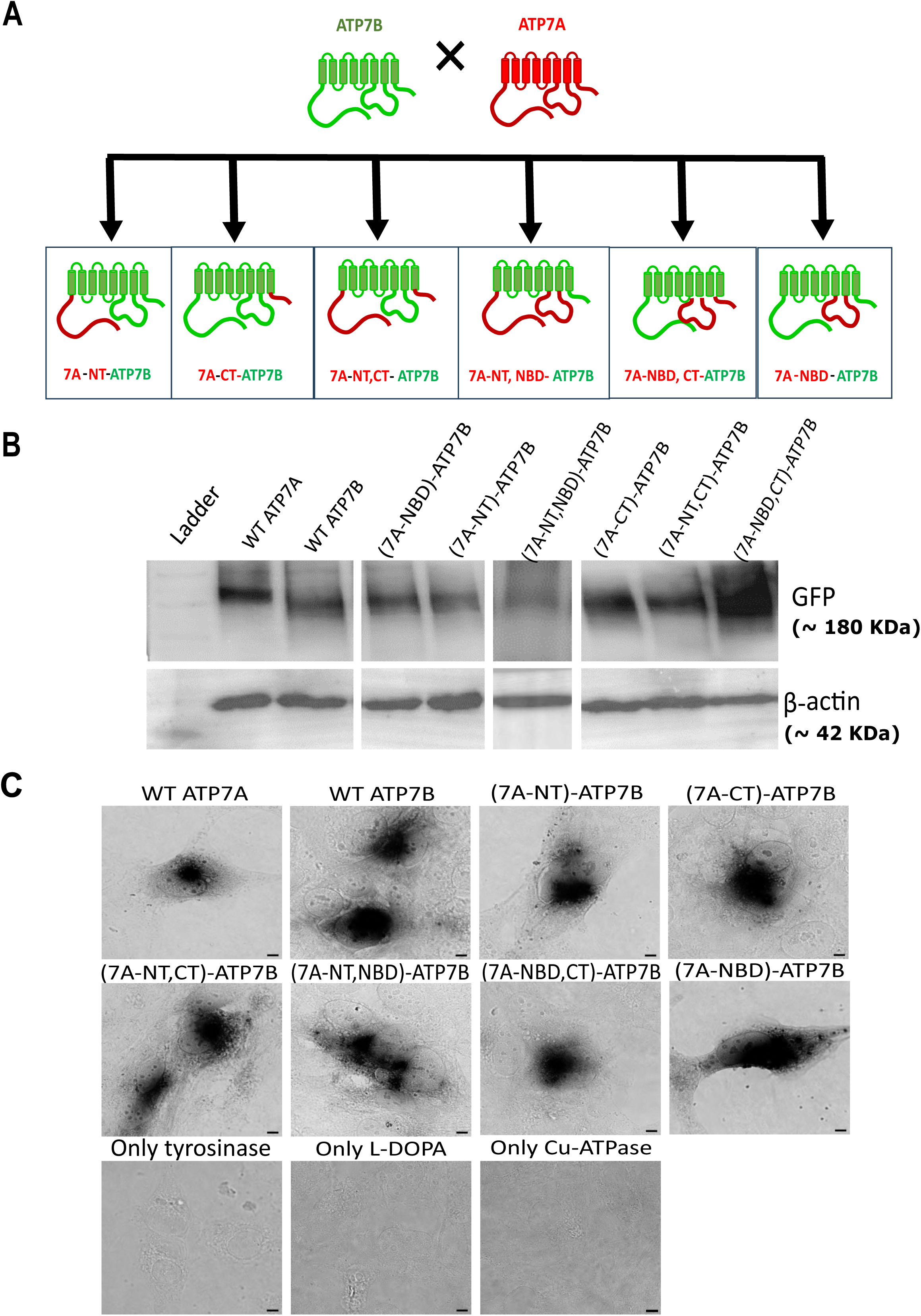
All chimeras are stably expressed and functionally active. (A) shows a schematic illustrating all the chimeras generated from ATP7A and ATP7B. (B) Immunoblot showing expression pattern of the six chimeras along with the wild type ATP7A (180kDa) and ATP7B (165kDa) proteins. β-actin (42kDa) is used as loading control (lower panel). All chimeras were stably expressed in HEK293T cells. (C) shows images of YS Menkes fibroblast cells transfected with the chimeras or wild type Cu-ATPase to check copper transport activity. All chimeric construct-transfected cells showed eumelanin pigmentation, indicating that they were functionally active, like the wild type Cu-ATPases. Scale bar: 10µm.

We then tested if these chimeric transporters could successfully pump copper ions across the membrane into the TGN lumen, by employing YS cell line - fibroblast cells isolated from a Menkes disease patient - which lack functional copper ATPases (Roy et al., 2020). Formation and precipitation of the eumelanin pigment in the cells were utilized as a read-out of Cu-ATPase activity that only occurs if the cuproenzyme tyrosinase receives copper as a co-factor in the TGN from a transiently-expressed copper transporter. All the six chimeric Cu-ATPases were observed to show copper-transport activity, as seen by the precipitation of eumelanin pigments in the cell, which was comparable with the wild-type Cu-ATPases, shown in **Fig. 6C**. Negative control YS cells that had been transfected with either just the Cu-ATPase construct or tyrosinase-encoding plasmid or cells that had not been transfected with any plasmid, but just treated with L-DOPA did not show any pigmentation.

### N-terminal, C-terminal and Nucleotide-binding domains of orthologous Cu-ATPases have undergone purifying selection

From the domain-substitution experiments, we noted that even though the chimeric Cu-ATPases were functionally active in terms of copper-transport activity, they exhibited unique localization and trafficking phenotypes. This suggested that though the copper transport function stays relatively conserved, the homologous transporters had undergone changes in their regulatory regions, making them adaptable for tissue-specific trafficking functions. This indication led us to further dissect the nature of the adaptive changes that have accumulated in these domains in the Cu-ATPases of the different life forms existing today. We looked at the Cu-ATPase domains of our distant kin to comprehend the extent of divergence between ATP7A and ATP7B. We retrieved the coding DNA sequences of the three cytosolic domains of 36 organisms spanning 13 phyla, ranging from unicellular bacteria to mammals. For ease of understanding, we clustered them into three taxonomic groups – a) Lower organisms’ group comprising unicellular organisms and lower invertebrates up to phylum Arthropoda, b) Intermediate organisms’ group consisting of phyla like mollusks, echinoderms, and invertebrate chordates (protochordates and cephalochordates) based on their evolutionary transition from prokaryotic and lower invertebrate metazoans to more complex organisms, and c) Higher organisms’ group comprising of vertebrates ranging from agnathans to mammals.

We then analyzed the mutation rates in the three cytosolic domains in all the different chosen taxa. As seen in **Fig. 7A**, NBD sequences of different taxa showed slightly higher overall mutation rate than the N-terminal and C-terminal domains. **Fig. S1A** shows a comparative analysis of the mutation rates in different evolutionary groups, while the table (**Fig. S1B**) shows the values obtained for each domain. In order to perceive the type of nucleotide substitutions that have occurred across the different taxa, we examined the transition/transversion (Ts/Tv) rates for all three domain sequences (**Fig. 7B)**. We found that there was a bias towards transition nucleotide-substitutions over transversions, in all the three domains analyzed, with the most pronounced bias seen in NBD, followed by the C-terminal and N-terminal domain. **Fig. S1C** depicts the differences in the (Ts/Tv) rates seen in different groups of organisms, across the domains and **Fig. S1D** shows the values obtained for each domain.

**Fig. 7.**
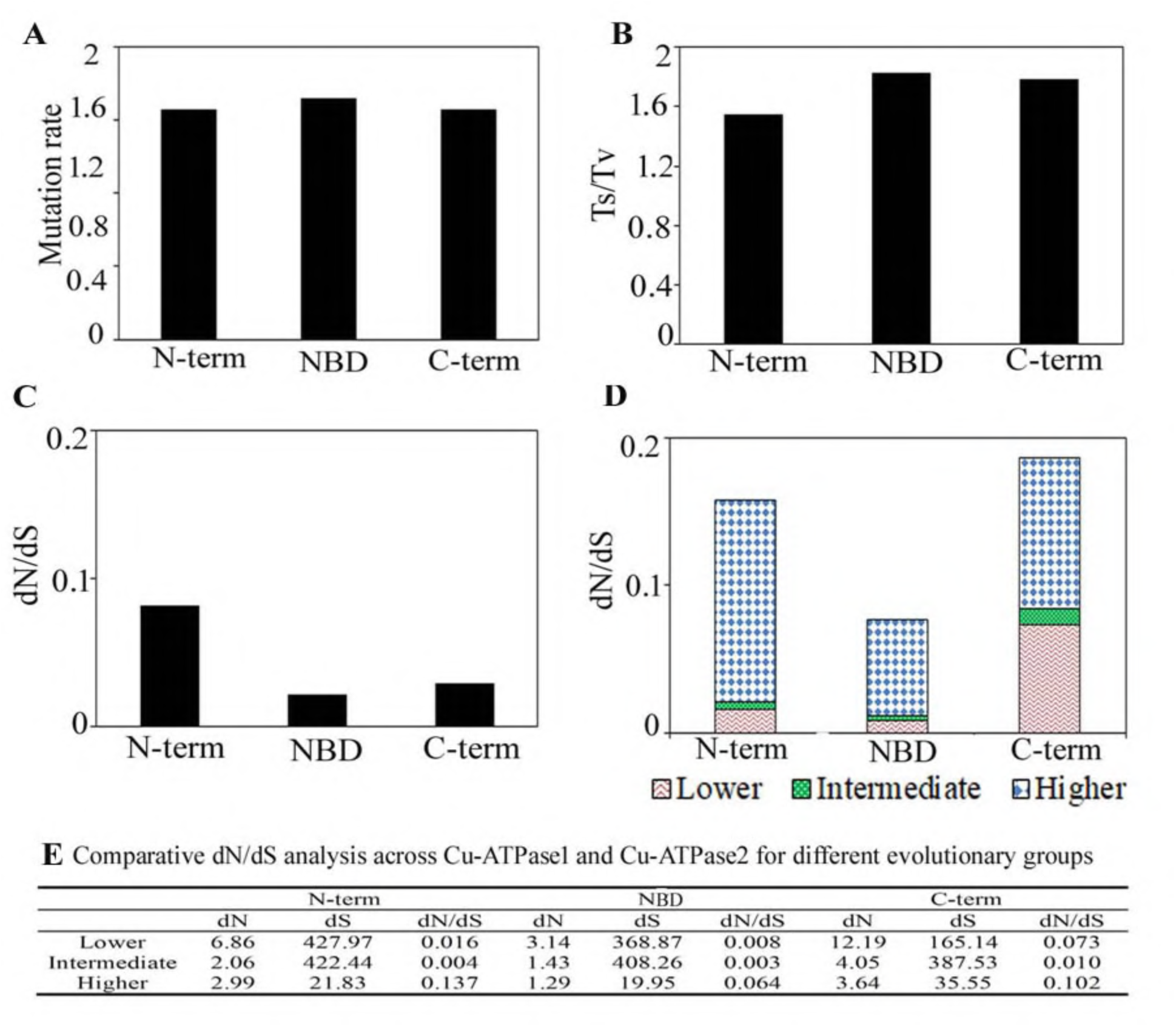
Evolutionary analysis of Cu-ATPases highlighting domain-specific mutation rates and selection pressures across all studied organisms. (A) Distribution of mutation rates across the N-terminal (N-term), Nucleotide Binding Domain (NBD), and C-terminal (C-term). (B)Transition/transversion (Ts/Tv) ratios for each domain, reflecting nucleotide substitution patterns. (C) Domain-wise dN/dS ratios (N-term, NBD, C-term) across all organisms. (D) dN/dS ratios categorized by taxonomic group: higher (blue), intermediate (green), and lower (red). (E) Comparative table of dN/dS values illustrating domain-specific selective constraints.

We next calculated the dN/dS ratio ̶ the balance between non-synonymous and synonymous substitutions in the three Cu-ATPase domains. We observed that across all species, all three domains had undergone purifying selection (dN/dS<1) as shown in **Fig. 7C**, with the NBD showing the lowest ratio, followed by the C-terminal and N-terminal domains, indicating that the NBD was the most conserved domain and C-terminal domain was the least. **Fig. 7D** shows the division of this evolutionary measure for different taxa examined. It was noted that the NBDs of the lower and intermediate taxa were subjected to stronger purifying selection as compared to higher organisms, hinting towards more relaxation in the selective pressures exerted on this domain in these organisms, possibly to facilitate regulatory changes. **Fig. 7E** shows the values obtained for the comparative dN/dS analysis for different evolutionary groups.

This explains, at least to an extent, why the NBD-substituted chimeric ATP7B constructs in our experiments failed to exit the TGN under high copper conditions; probably these regions of the proteins had acquired changes in their regulatory regions that made them irreplaceable in terms of trafficking.

### N-terminal and Nucleotide-binding domain trees show similar evolutionary trends

We analyzed the CDS sequences of the Cu-ATPase domains and constructed phylogenetic trees using IQ-TREE (v2.2.0) with the Maximum Likelihood (ML) method to visualize the evolutionary patterns of these conserved domains. **Fig. 8A** and **8B** shows the phylogenetic trees obtained for N-terminal domain and Nucleotide Binding Domain (NBD) respectively. Notably, both these trees exhibited comparable clustering between the three taxonomic groups, especially the differences between the higher organisms and lower organisms. The C-terminal domain, on the other hand, showed considerable inter-mixing of different taxa as seen in **Fig. 8C**. Among vertebrates, the ATP7B C-terminal sequences showed closer relationships with each other than those of ATP7A. These tree patterns underline the evolutionary forces acting on various domains of Cu-ATPases reflecting their functional adaptations in organisms across evolution.

**Fig. 8.**
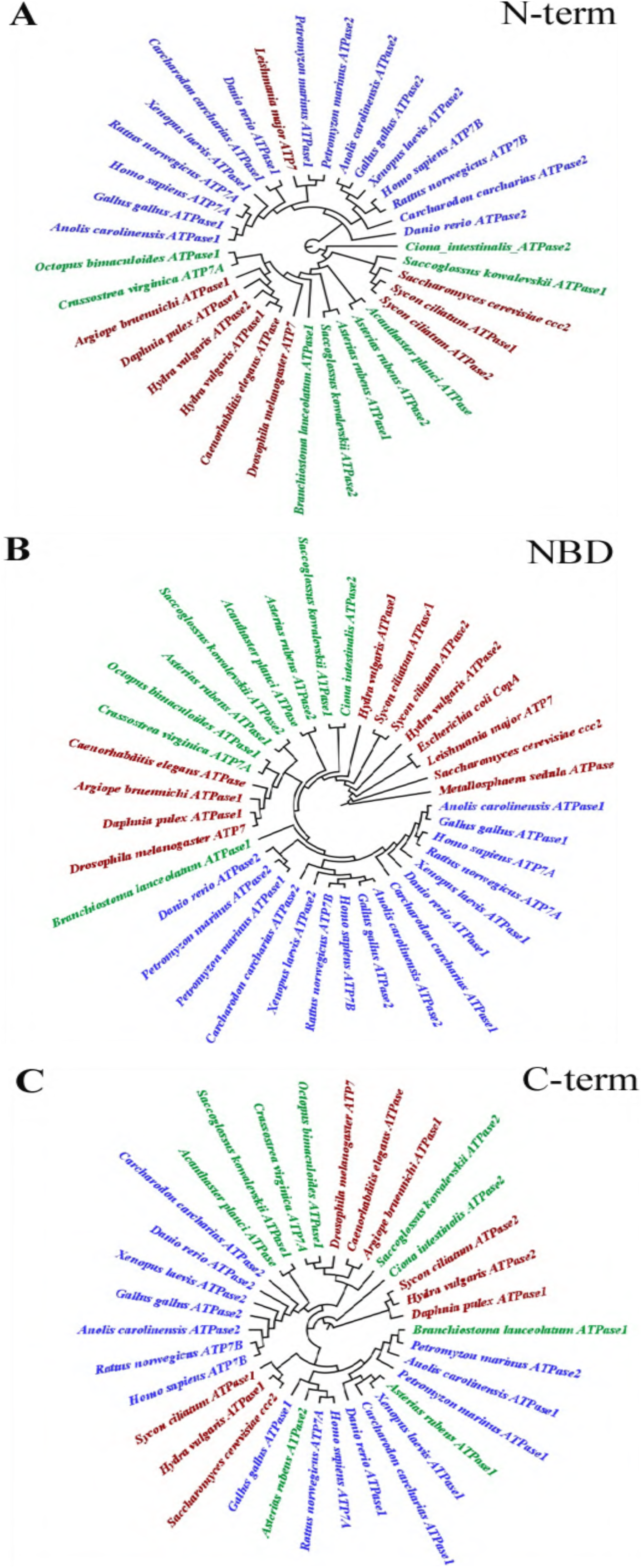
Phylogenetic trees illustrating the evolutionary relationships of Cu-ATPases, constructed using the Maximum Likelihood (ML) method. **Taxa are color-coded by evolutionary hierarchy –** higher organisms (from Agnatha to mammals) are shown in blue; intermediate taxa (including Mollusca, Echinodermata, Hemichordata, and Cephalochordata) are shown in green; and lower organisms (Archaea, Bacteria, Protista, Porifera, Cnidaria, Nematoda, and Arthropoda) are depicted in red. Panels B–D present domain-specific phylogenetic trees of Cu-ATPases: (B) N-terminal domain (N-term), (C) Nucleotide Binding Domain (NBD), and (D) C-terminal domain (C-term).

**Fig. 9.**
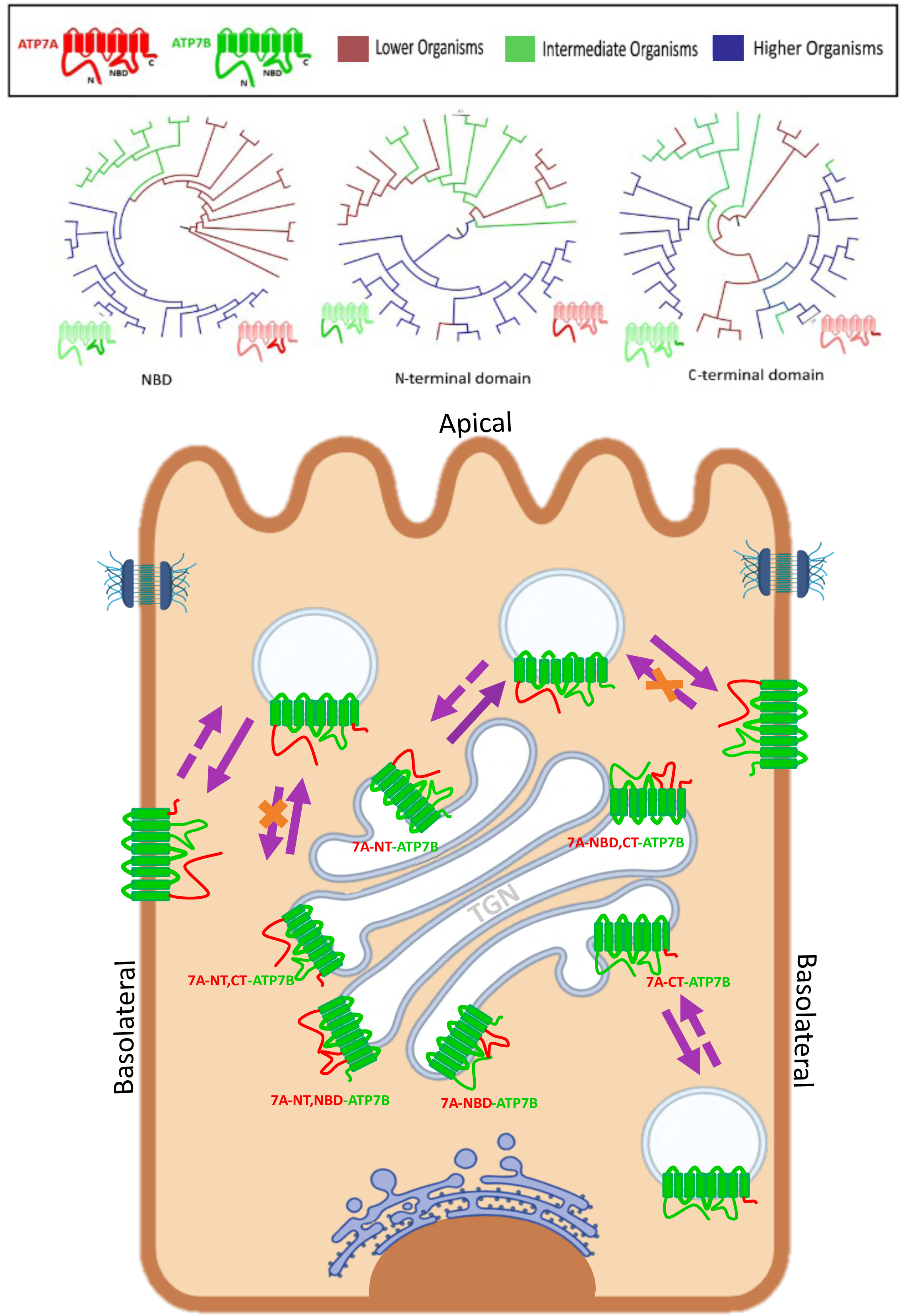
Graphical abstract summarizing the evolutionary and experimental part of the study. Upper panel depicts the domain-wise phylogenetic trees showing evolutionary relationships between different organisms. Lower panel depicts the varied trafficking phenotypes of the domain-swapped ATP7B chimeras, in polarized cells. Notably, when the N-terminal domain of ATP7B was substituted with that of ATP7A (7A-NT-ATP7B), the protein constitutively trafficked to the basolateral membrane in polarized cells, but additionally replacing the NBD led to the TGN-retention of this doubly-substituted chimera. Substituting just the C-terminal domain did not affect the Cu-dependent vesicularization or recycling of the chimeric ATP7B to the TGN. All NBD-substituted chimeras, however, failed to exhibit Cu-dependent trafficking.

## Discussion

Copper-transporters are conserved polytopic P-type ATPases, crucial for maintaining optimal concentrations of copper in all organisms. Cu-ATPase homologs in vertebrates - ATP7A and ATP7B - show tissue-specific expression and distinct trafficking polarities. They traffic between the TGN and the apical (ATP7B) or the basolateral (ATP7A) membranes in polarized epithelial cells, depending on the intracellular copper levels. In the current study, we set out to delineate the roles that the homologous cytosolic domains constituting these Cu-ATPases play in mediating their trafficking and copper-transport activity. We adopted a domain-substitution approach for the same since the two homologs diverged from a single Cu-ATPase, by reasoning that swapping their conserved domains would provide insights into interdomain interactions and conformations of the Cu-ATPases.

The large Cu-sensing N-terminal domain, Nucleotide-binding and Phosphorylation domains (NBD) and the C-terminal domain were considered for our study as they have been reported to be important for trafficking regulation, besides copper-transport. We found that the ATP7B chimeric constructs with N-terminal domain/NBD/C-terminal domain(s) of ATP7A were all functionally active. Cryo-EM ATP7B structures have led to the conclusions that interactions between metal-binding subdomain 6 of the ATP7B N-terminal with the A-domain, and that of A-domain with P-domain, accompanied by a change in the motion of the A-domain subsequently exposes the transmembrane Cu-bound site to the lumen to facilitate copper efflux (Yang et al., 2023). Such information with regards to ATP7A, however, is lacking. Domain-substitution of ATP7B with ATP7A indicates that the regions/motifs involved in this complex process are conserved in both the Cu-ATPase homologs.

All the chimeric constructs showed diverse localization and trafficking phenotypes. We found that substituting the N-terminal domain of ATP7A in ATP7B, made the protein lose predominant steady-state TGN-localization unlike wild type ATP7A and ATP7B at low copper or basal condition. Similar phenotype was seen with double-substitution of ATP7B N-terminal and C-terminal domains. Both these chimeras trafficked to the basolateral membrane. This observation corroborates the findings by Braiterman et al. who also implicated the ATP7B N-terminal domain (1-63 amino acid region) in copper-dependent TGN-localization and apical targeting (Braiterman et al., 2009). The Wilson disease-causing N41S ATP7B mutation within the nine amino acid stretch F^37^AFDNVGYE^45^ implicated in apical targeting was also mistargeted to the basolateral membrane (Braiterman et al., 2009; Ruturaj et al., 2024). Interestingly substituting the ATP7B C-terminal domain with that of ATP7A not only did not affect the steady-state TGN localization of the chimera in polarized cells, it showed Cu-responsive trafficking into post-TGN compartments, although it did not reach either plasma membrane domains in polarized cells. But in unpolarized cells, this chimera was trafficked to the cell membrane to an extent, indicating that the ATP7A C-terminal domain motifs (dileucine,^1487^LL^1488^ or ^1497^DTAL^1500^) could be recognized at this stage by trafficking regulators, however owing to a shift in the trafficking machinery that occurs during polarization of epithelial cells, this chimera now can only exit the TGN to move into post-TGN compartments. Braiterman et al. had performed trafficking studies in polarized WIF-B cells (hepatocytes) with ATP7B in which ATP7A C-terminal had been substituted; intriguingly, they had observed this chimera to traffic to both apical and basolateral membranes in those cells (Braiterman et al., 2011). This may be due to differences in the trafficking machinery of different cell types. It is worthwhile noting that the chimeras with a cis-ATP7B-N-term (in combination with other ATP7A domains) could not traffic apically even in the presence of high copper; similar was the case with chimeras with cis-C-terminal domains. These observations indicate that both N-term and C-term motifs are necessary, but not sufficient for apical targeting of ATP7B, collectively, making a strong case for the necessity of inter-domain interactions for the correct trafficking of the protein. The ATP7B C-term trileucine motif, L^1454^-L^1456^ has been reported to be essential for retrograde trafficking, however, the chimera with C-term of ATP7B replaced with that of ATP7A could also traffic back to the TGN, indicating that the retrograde trafficking machinery was conserved among the homologous C-terminal domains, most probably due to the presence of the dileucine motif in both. Moreover, the presence of the ATP7A C-terminal domain in the chimeric protein containing ATP7A N-terminal domain exhibited retrograde trafficking, though incompletely, from the basolateral membrane on copper chelation, signifying the role of the C-terminus (in presence of N-terminus) in sorting of the protein in vesicles.

The most remarkable phenotype revealed from our study was by that of the (7A-NT, NBD)-ATP7B chimera. While 7A-NT-ATP7B trafficked to the basolateral membrane even at low copper conditions, additionally replacing the ATP7B NBD with that of ATP7A, made the resultant double-substituted chimera to localize at the TGN. This hints towards a TGN-stabilizing interaction between ATP7A N-terminal domain and NBD, as also hypothesized to occur under low copper conditions by Hasan & Gupta et al. (Hasan & Gupta, 2012). When the NBD was substituted in ATP7B, singly or along with N-terminal/C-terminal domains, the chimeras were localized at the TGN mostly, at even high copper conditions, in MDCK. It has been postulated by Hasan et al. that the for high-copper-induced TGN-exit, ATP7B loses interaction between its copper-bound N-terminal domain and the N-domain (part of NBD). Unfolded loops in N-domain have been hypothesized to be significant for their trafficking regulation (Dmitriev et al., 2006; Yatsunyk & Rosenzweig, 2007). We hypothesize that the chimeric NBD-substituted Cu-ATPases, although functionally active, fail to adopt this open trafficking-ready conformation and end up masking the key trafficking motifs. Previous studies from our group identified regulatory partners of ATP7A and ATP7B by mass-spectrometry studies, that included clathrin-dependent Adaptor Protein-1 complex, which was found to be essential for correct sorting of both copper-ATPases in polarized MDCK (Ruturaj et al., 2024). It is also possible that these chimeric proteins end up being sorted into an ‘incorrect’ TGN-sub-domain.

To further interpret the observations from domain-substitution studies, we studied the selective pressures shaping the Cu-ATPase cytosolic domains of our interest. Although there have been studies on the evolution of Cu-ATPase proteins in various organisms (Fodor et al., 2023; Migocka, 2015; Gupta & Lutsenko, 2012), our domain-specific phylogenetic studies revealed new insights that supported the experimental results of our study. The NBD was found to be highly conserved relative to the other tested domains, highlighting its essential roles in ATP binding and hydrolysis as also shown by previous studies (Tsivkovskii et al., 2001). The N-terminal domain, crucial for copper sensing, exhibited considerable variability across taxa, indicating adaptive mechanisms in response to fluctuating copper availability in diverse environments (Lutsenko et al., 1997), similar to the evolutionary pattern of the extracellular amino-terminal domain of the high-affinity copper importer CTR1 crucial for copper sequestration from the environment (Kar et al., 2022). However, the phylogenetic trees of both N-terminal domain and NBD showed a similar pattern with close clustering between vertebrate Cu-ATPases and closer relationships between human homologs. This was unlike the pattern shown by the C-terminal domain sequences, which remarkably showed no apparent clustering between the evolutionary groups. The consistently low dN/dS ratios across all three Cu-ATPase domains, among all taxa, reflected purifying selection in a bid to preserve the functional integrity of the proteins. The N-terminal and C-terminal domains exhibited slightly higher dN/dS values, indicating lesser stringent selection acting on these regulatory domains than on the NBD. The Ts/Tv rates revealed a pronounced transition bias, especially in the NBD, reflecting its essential functional role across all taxonomic groups.

To summarize, the functional interplay between the different cytosolic domains– N-terminal, NBD, and the C-terminal domain is crucial for TGN localization and exit in response to copper. Substituting N-terminus, NBD, or C-terminus of ATP7B, either singly or in pairs, with the corresponding domain(s) of its homolog ATP7A, gave rise to chimeric proteins that were functionally active but showed altered trafficking regulation. Overall, our studies suggest that the three key cytosolic domains in the chordate Cu-ATPase homologs have accumulated changes through the course of selection that render them indispensable for correct copper-induced trafficking.

### Methodology

#### Plasmids and Antibodies

eGFP-ATP7B construct was available in the lab previously (Das et al., 2020). eGFP-ATP7A construct was made on the existing mKO2-HA-ATP7A (Ruturaj et al., 2024) plasmid using NEBuilder HiFi DNA Assembly (NEB #E2621). Chimeric constructs were also prepared by NEBuilder HiFi DNA Assembly according to manufacturer’s protocol. The entire chimera cloning is schematically presented in **Fig. S2A** Primer details are in **Fig. S2B** The domains of the two homologous Copper ATPases that were replaced, encompasses the following residues: For hATP7A, N-terminal domain: 1-645 amino acids; NBD loop (N-domain+P-domain): 1004–1348; C-terminal domain: 1398-1492. For hATP7B, N-terminal domain: 1-653; NBD loop (N-domain+P-domain): 995-1322; C-terminal domain:1372-1465. The tyrosinase plasmid pcTYR was a kind gift from Prof. Svetlana Lutsenko, Johns Hopkins University, USA. Plasmid isolation was done using Macherey Nagel plasmid isolation kit (#MN740490).

Following are the antibodies that were used for the study: rabbit anti-Golgin97 (CST # 13192); 1:500, mouse anti-ATP1A1 (Abcam #ab7671); 1:500, mouse anti-gp135 (DSHB #3F2/D8); 1:400, rabbit anti-GFP (BioBharati#BB-AB-0065);1:5000, mouse Beta actin (Proteintech #66009-1-Ig); 1:25,000, Phalloidin-iFluor 405 Reagent (Abcam #ab176752);1:1200, anti-mouse Alexa 555 (Invitrogen #A32773); 1:800, anti-rabbit Alexa 568 (Invitrogen #A11011); 1:800, anti-rabbit Alexa 647 (Invitrogen #A32733); 1:800, anti-rabbit HRP conjugated (CST #7074S); 1:5000, anti-mouse HRP (CST #7076, 1:6000;)

#### Cell lines and Cell culture

MDCK cells (a kind gift from Prof. Enrique Rodriguez-Boulan’s laboratory, Weill Cornell Medical College, USA) were grown and maintained in media consisting of Dulbecco’s modified Eagle’s medium (DMEM Gibco #11995065), supplemented with 10% FBS (Gibco #26140079) and 1X PenStrep (Gibco #15140122) at 37 ͦC and 5% CO2. For polarization, 3 x 10^5^ cells were plated in 0.4μm, 12mm transwell inserts (Corning #3401) and grown for 4-5 days. For transfection, electroporation was performed using Nucleofector 2b device (Lonza #AAB-1001) and Amaxa Kit V (Lonza #VCA-1003), Program T-023. HEK293T (a kind gift from Prof. Arindam Mukherjee’s laboratory, Indian Institute of Science Education and Research Kolkata, India) and YS Menkes patient’s fibroblast cell line (a kind gift from Prof. Svetlana Lutsenko’s laboratory, Johns Hopkins University, USA) were cultured in DMEM supplemented with 10% FBS (Gibco # 10270-106), 1X PenStrep. For transfection, JetPRIME transfection reagent (#114-07; Polyplus) was used as per the manufacturer’s protocol.

For copper treatment, 75µM of CuCl2 was used for 1 hr. For mimicking copper-deprived conditions, cells were treated with 100µM BCS for 2hrs. Both the reagents were prepared in autoclaved deionized water.

#### Immunofluorescence and microscopy

After desired treatments, cells were washed twice with ice-cold PBS, then fixed with 2% PFA for 20 mins. Following a PBS rinse, cells were incubated in 50mM ammonium chloride solution in PBS. Next, the cells were washed with PBS, and blocking was performed, along with permeabilization, in 1% Bovine Serum Albumin (BSA, SRL #85171) in PBSS (0.075% saponin in PBS) for 20 mins. Primary antibody incubation was performed for 2 hrs at RT followed by three PBSS washes. This was followed by a 1 hour incubation with the respective secondary antibodies. The cells were then washed thrice with PBSS and PBS, respectively. The membrane was mounted on a glass slide with ProLong Gold Antifade Reagent (CST #9071S). All images were acquired with Leica SP8 confocal platform using oil immersion 63x objective (NA 1.4) and deconvolution was done using Leica Lightning software. All the images were captured at z-interval of 0.25 μm.

#### Tyrosinase assay

YS cells grown on glass coverslips were transfected with either 0.1μg of pcTYR or 0.25 μg of plasmid encoding Cu-ATPase (wt or chimera) or both expression plasmids using JetPrime transfection reagent.

Eighteen hours later, cells were washed twice in PBS and fixed for 30 s in cold acetone-methanol (1:1). These fixation conditions do not interfere with tyrosinase activity. The cells were then incubated for 4 hrs at 37 °C in 0.1 M Na-phosphate buffer (pH 6.8) containing (0.4mg/ml) levo-3,4-dihydroxy-Lphenylalanine (L-DOPA) freshly prepared, in dark. Coverslips were mounted on slides with Prolong Gold mounting media without DAPI, and formation of the black L-DOPA chrome pigment was detected by bright-field microscopy.

#### Image Analysis

All images were processed and analyzed using FIJI software. For colocalization calculation, the JaCoP plugin was used. The Macro code for the analysis is available at https://github.com/saps018/7A-7B. Manders’ Colocalization Coefficient was calculated from manually drawn ROIs. Non-parametric tests for unpaired datasets (non-parametric T-test) was performed for all the samples; \**p* < 0.05; \*\**p* < 0.01; ***p<0.001; \*\*\*\**p* < 0.0001; ns, not significant. Graphs were plotted using Origin 2018 software and statistical significance was calculated using Microsoft Excel (2013). For interpretation of violin plots, Lower Quartile (Q1 or 25th Percentile) – The bottom dashed line represents the first quartile, meaning 25% of the data falls below this value; Median (Q2 or 50th Percentile) – The middle dashed line represents the median, which is the central value of the data distribution; Upper Quartile (Q3 or 75th Percentile) – The top dashed line represents the third quartile, meaning 75% of the data falls below this value.

#### Immunoblot analysis

HEK293T cells were transfected with either the wild type or chimeric copper transporter-encoding construct using JetPrime reagent, according to manufacturer’s instructions and after 24 hours, cells were pelleted down and pellet was resuspended in 100 μL of RIPA lysis buffer (10 mM Tris-Cl pH 8.0, 0.5 mM EGTA, 1 mM EDTA, 1.0% Triton X-100, 0.1% sodium deoxycholate, 0.1% SDS, 140 mM NaCl, 1X protease inhibitor cocktail, 1 mM phenylmethylsulfonyl fluoride [PMSF]) and incubated on ice for 20 min. The cells were then sonicated with a probe sonicator (six pulses, 5 s, and 100 mA). Quantitation of protein was done using Bradford reagent and the protein samples were mixed with 4X NuPAGE loading buffer (#NP0007; Invitrogen) to a final concentration of 1X and electrophoresed on 8% SDS polyacrylamide gel. The protein bands were then transferred onto PVDF membrane (#IPVH00010 Millipore) by semi-dry transfer apparatus. This was followed by blocking with 3% BSA in TBST for 2hrs and antibody incubation step. The membrane was probed with primary antibody against GFP or beta-actin at 4°C overnight on a rocker, followed by two hours of HRP-conjugated secondary antibody incubation. Then, the membranes were washed with TBST and TBS before visualizing the signal with ECL developer (Bio-Rad #1705062) using ChemiDoc (Bio-Rad).

#### Software and programme

Programme written in Python-language is available at GitHub in the following link. (https://github.com/AAYATTI/Evolution). Further queries regarding computer programme may contact AMG

#### Sequence retrieval and alignment

All CDS and protein sequences used in the study were retrieved from NCBI database (https://www.ncbi.nlm.nih.gov/). Domain annotations of the required sequences were fetched from Uniprot (https://www.uniprot.org/) and DeepTMHMM (Hallgren et al., 2022). Multiple sequence alignment was performed using **MAFFT** (v7.505) (Katoh and Standley, 2013), a widely used tool for efficient and accurate alignment of nucleotide and protein sequences. The input dataset consisted of curated sequences in FASTA format, ensuring no gaps, ambiguous residues, or redundancies. The alignment strategy was automatically determined by MAFFT based on the dataset’s size and complexity to ensure optimal alignment quality. The resulting aligned sequences were saved in FASTA format for subsequent analyses. Pairwise sequence alignment of the vertebrate copper-ATPases were done using EMBOSS Needle server (Madeira et al., 2024).

#### Phylogenetic Tree Construction

Phylogenetic tree construction was carried out using **IQ-TREE** (v2.2.0) (Minh et al., 2020) with the Maximum Likelihood (ML) method. ModelFinder (Kalyaanamoorthy et al., 2017) was employed to systematically test and select the best-fit substitution model based on the Bayesian Information Criterion (BIC), ensuring an optimal evolutionary framework for tree construction. Ultrametric bootstrap analysis was performed with 1,000 replicates to assess the statistical robustness of the tree topology, with branch support values integrated into the final tree. Computational efficiency was optimized by automatically adjusting the number of processing threads based on the available system resources. The resulting tree was output in Newick format and visualized using **iTOL** (Interactive Tree of Life) (Letunic et al., 2021) for interpretation and presentation. For the C-terminal domain tree, unicellular organisms could not be included due to the short lengths of this domain in these organisms.

#### Calculation of dN/dS Ratio for Selective Pressure Analysis

The nonsynonymous to synonymous substitution rate ratio (dN/dS) was calculated using **CODEML**, a program within the **PAML** (Phylogenetic Analysis by Maximum Likelihood) package (Yang, 2007),to assess the selective pressures acting on the sequences. The analysis was performed using a codon-based alignment of nucleotide sequences, ensuring that proper reading frames were maintained and that no stop codons or frameshift errors were present. A phylogenetic tree, constructed in Newick format, was used as input along with the aligned sequences. The dN/dS ratio, denoted as ω, was calculated by CODEML, providing estimates for dN, dS, and ω values across the dataset. A ratio of ω > 1 indicates positive selection, ω = 1 suggests neutral evolution, and ω < 1 points to purifying selection. In some cases, additional models, such as branch-specific or site-specific models, were tested to explore variability in selective pressures across different branches or sites of the phylogeny, offering a more nuanced understanding of evolutionary dynamics in the dataset.

#### Estimation of Transition/Transversion (Ts/Tv) Ratio

The Transition/Transversion (Ts/Tv) ratio were estimated using **IQ-TREE** (v2.2.0) (Minh et al., 2020) as part of the substitution model fitting process. IQ-TREE reflects the relative rate of transitions (purine to purine or pyrimidine to pyrimidine) compared to transversions (purine to pyrimidine or vice versa). The Ts/Tv ratio provided insights into the nucleotide mutation patterns. Higher Ts/Tv ratio values indicate a greater propensity for transitions, a characteristic commonly observed in certain evolutionary processes.

#### Determination of mutation rate

The mutation rate was calculated using a customized Python script that determines the ratio of parsimony informative sites to the total number of nucleotide sites in the sequence alignment. The values for parsimony informative sites and total sites were obtained from the IQ-TREE analysis. Parsimony informative sites are defined as positions in the alignment where at least two distinct nucleotides are observed, which are essential for phylogenetic analysis, while total sites refer to all nucleotide positions in the alignment. This ratio provides a quantitative measure of mutation frequency, offering insights into the extent of nucleotide variation across the dataset.

## Acknowledgement

This work was supported by DBT-Wellcome Trust India Alliance Fellowship (IA/I/16/1/502369) and Core Research Grant (CRG/2021/002150) from SERB, Department of Science and Technology (DST), Government of India, STARS-2 Grant (2023-0210) from Ministry of Education, Govt. of India and IISER K intramural funding to A.G. S.M.J. is supported by pre-doctoral fellowship from University Grants Commission, India. S.M. and M.P. are supported by pre-doctoral fellowship from Council of Scientific and Industrial Research, India. A.M.G. is supported by National Postdoctoral Fellowship from SERB, Department of Science and Technology (DST), Government of India.

## Author contributions

Conceptualization: A.G. and A.M.G. Methodology: A.G., S.M, S.M.J, M.P. and A.M.G; Software: A.M.G.; Formal analysis: A.G., S.M, S.M.J and A.M.G; Resources: A.G.; Data curation and analysis: A.G., A.M.G and S.M.J; Writing - original draft: A.G., A.M.G and S.M.J.; Project administration: A.G.; Funding acquisition: A.G.

## Competing interests

The authors declare no competing interests.

**The manuscript does not require ethics committee approval at the institution.**

## Data availability statement

The authors confirm that the data supporting the findings of this study are available within the article and/or its supplementary materials.

---

**Fig. S1:**
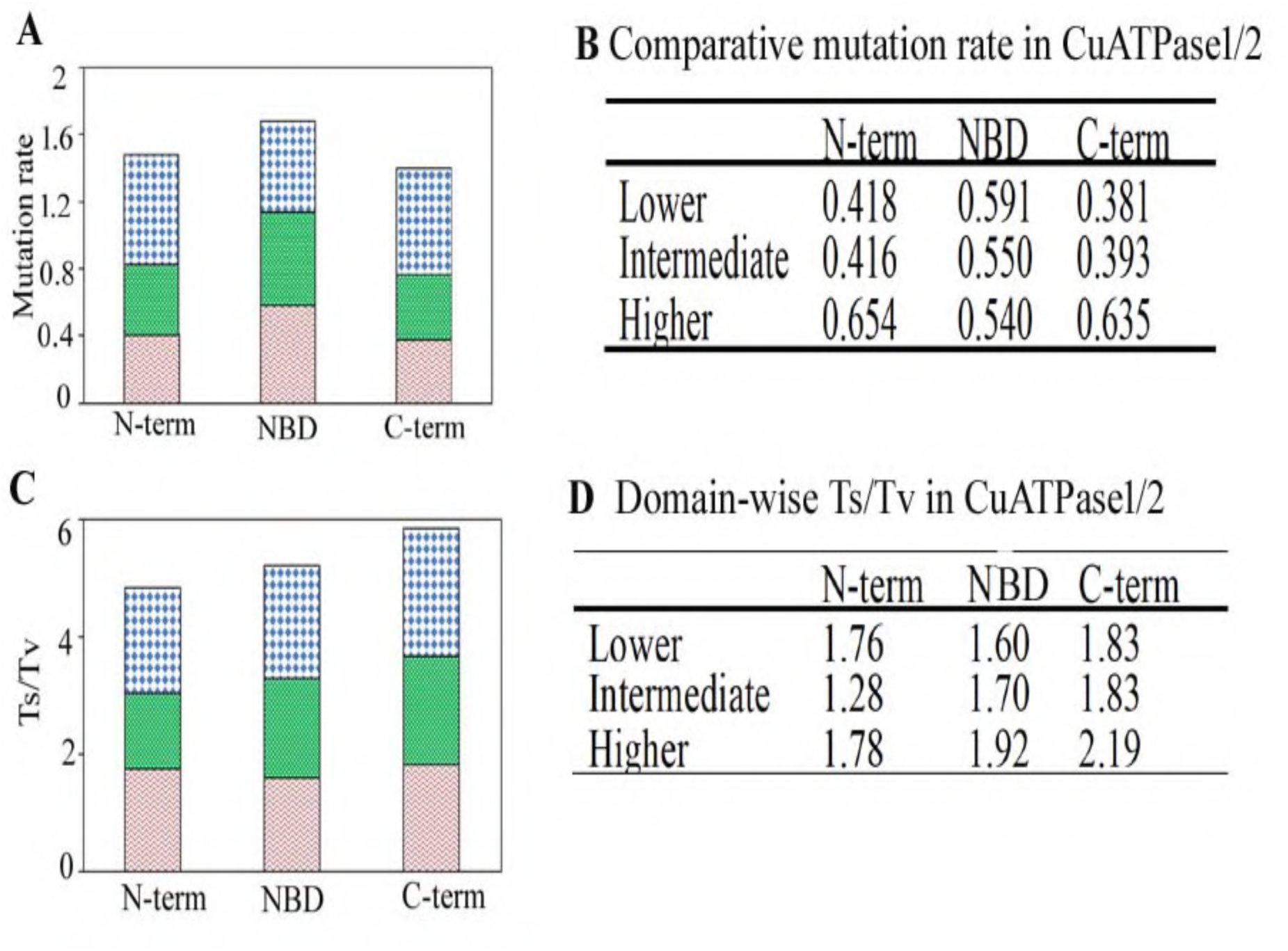
Evolutionary analysis of Cu-ATPases, illustrating domain-specific mutation rates and transition/transversion (Ts/Tv) ratios across taxa. Different evolutionary parameters were checked for the Cu-ATPase domains of higher (blue), intermediate (green), and lower (red) organisms. (A) Distribution of mutation rates for the N-terminal (N-term), Nucleotide Binding Domain (NBD), and C-terminal (C-term) across different taxonomic groups. (B) Table summarizing mutation rates for each domain and taxonomic category. (C) Ts/Tv ratios for the three domains, reflecting nucleotide substitution patterns. (D) Table presenting domain-wise Ts/Tv values across taxa.

**Fig. S2:**
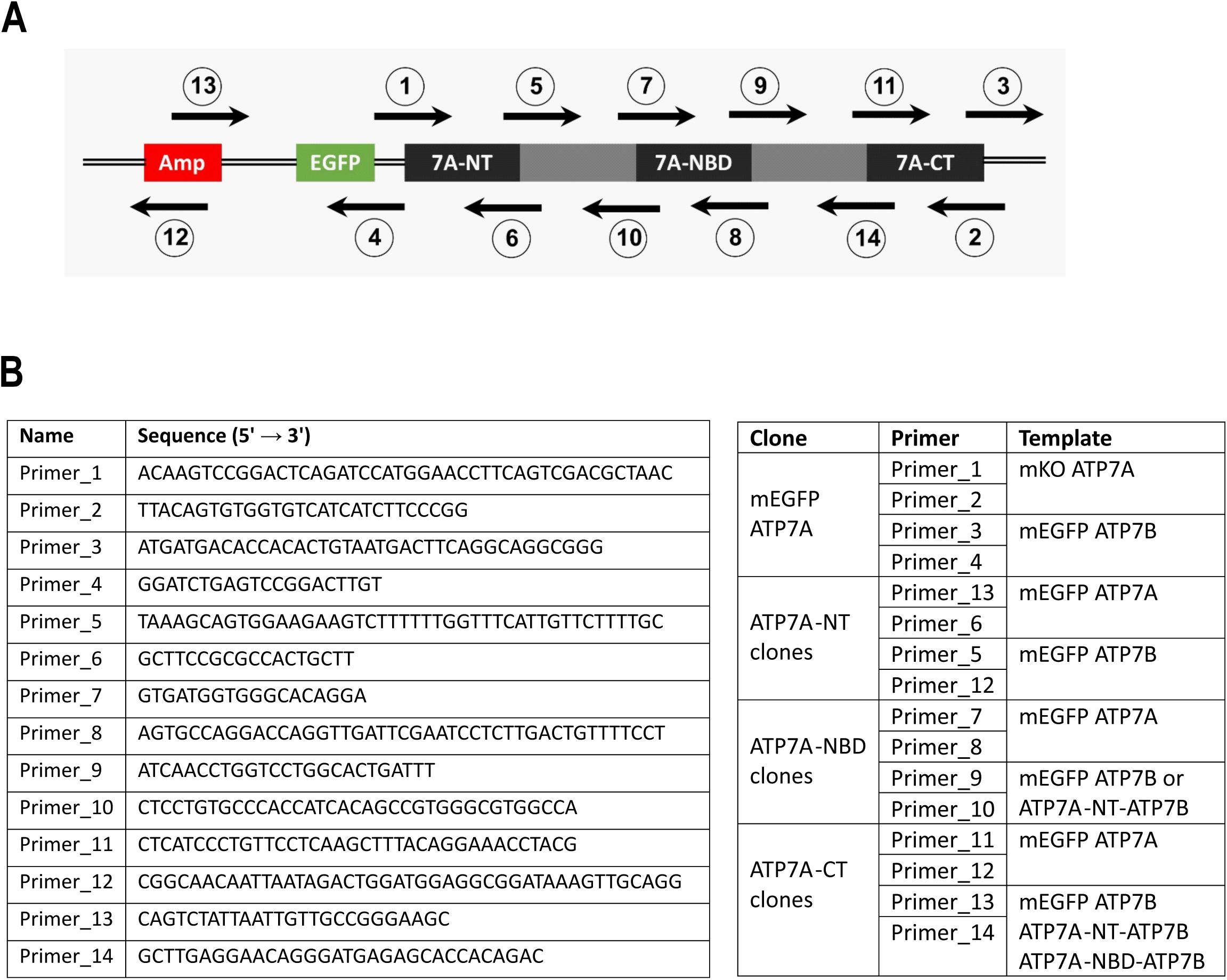
Chimera cloning schematic and primer details. (A) Schematic showing location of labelled primers that were used for generating chimeras by Hi-Fi cloning. (B) Table showing primer sequences used for chimera cloning.

